# Stochastic bond dynamics facilitates alignment of malaria parasite at erythrocyte membrane upon invasion

**DOI:** 10.1101/2020.03.01.971986

**Authors:** Sebastian Hillringhaus, Anil K. Dasanna, Gerhard Gompper, Dmitry A. Fedosov

## Abstract

Malaria parasites invade healthy red blood cells (RBCs) during the blood stage of the disease. Even though parasites initially adhere to RBCs with a random orientation, they need to align their apex toward the membrane in order to start the invasion process. Using hydrodynamic simulations of a RBC and parasite, where both interact through discrete stochastic bonds, we show that parasite alignment is governed by the combination of RBC membrane deformability and dynamics of adhesion bonds. The stochastic nature of bond-based interactions facilitates a diffusive-like re-orientation of the parasite at the RBC membrane, while RBC deformation aids in the establishment of apex-membrane contact through partial parasite wrapping by the membrane. This bond-based model for parasite adhesion quantitatively captures alignment times measured experimentally and demonstrates that alignment times increase drastically with increasing rigidity of the RBC membrane. Our results suggest that the alignment process is mediated simply by passive parasite adhesion.

## I. INTRODUCTION

Malaria is a dangerous mosquito-borne disease which kills nearly 0.5 million of people every year [1]. It is caused by a protozoan parasite of the genus Plasmodium and proceeds in several stages [2–4]. After about 10 days from the initial infection through a mosquito bite, an infected liver releases a large number of merozoites, egg-shaped parasites with a typical size of 1 – 2*μ*m [5, 6], into the blood stream. The blood stage of malaria infection is a clinically relevant stage, where merozoites invade healthy red blood cells (RBCs) and multiply inside by utilizing the RBC internal resources. This intra-erythrocytic development is essential for merozoites to be hidden from the immune system and avoid clearance. After about 48 hours post RBC invasion, infected RBCs are ruptured and new merozoites are released into the blood stream to repeat this reproduction cycle. Thus, RBC invasion by merozoites is crucial not only for parasite survival, but also for further multiplication.

RBC invasion by merozoites is preceded by three key events: (*i*) initial attachment, (*ii*) re-orientation or alignment of the parasite such that its apex is facing the RBC membrane, and (*iii*) formation of a tight junction [7]. The apex contains all required machinery to invade RBCs after the tight junction is formed [8]. At physiological hematocrit levels with a volume fraction of RBCs close to 40%, initial attachment of merozoites can be considered almost immediate after their egress from infected RBCs. However, the initial attachment has a random parasite orientation, which rarely provides direct alignment of the apex toward the membrane required to start the invasion. This implies that the parasite alignment is an extremely crucial step for successful invasion, which needs to be completed within a couple of minutes, as after this time period merozoites generally lose their ability to invade RBCs [9]. To facilitate parasite alignment, merozoites contain a surface coat of proteins, mainly GPI-anchored, which can bind to the RBC membrane [5, 10, 11]. However, one of the main difficulties in the investigation of RBC-parasite interactions is that exact receptor-ligand bindings remain largely unknown. Electron microscopy images [5] of merozoites adhered to a RBC suggest that along with short surface-coat filaments of length ≃ 20 nm, there exist much longer filaments of lengths up to 150 nm, which may play an important role in early stages of merozoite adhesion to the RBC membrane. Furthermore, these long filaments have a much lower density than short binding filaments. Even though adhesion kinetics of such bonds remain unknown, recent optical tweezers experiments [9] indicate the adhesion force of spent merozoites to the RBC membrane to be within the range of 10 to 40 pN.

Another important aspect during merozoite alignment is the deformation of the RBC membrane. Dynamic membrane deformations of various magnitudes are often observed [12–15] and are thought to aid in the alignment process [16, 17]. Recent live-cell imaging experiments show a positive correlation between RBC deformations and eventual merozoite alignment [16]. A deformation score on the scale from 0 to 3 with increasing membrane deformation has been introduced. Most merozoites that successfully invade RBCs induce a deformation score of either 2 or 3, while for deformation scores of 0 or 1 the invasion success is much less frequent [16]. Furthermore, these experiments lead to an estimate of an average alignment time of about 16 s [16]. A recent simulation study by [17], with RBC-parasite adhesion modeled by a homogeneous interaction potential, has confirmed the importance of membrane deformations, which facilitate parasite alignment through its partial wrapping by the membrane. However, this model shows static (not dynamic) membrane deformations and leads to average alignment times of less than 1 s, indicating that an essential aspect of the alignment process has not been captured. Another speculation is that dynamic membrane deformations are induced actively by merozoites through changing locally the concentration of Ca^+^ ions [18, 19]. This proposition has been confronted by recent experiments [20], which show that calcium release by parasite starts only at the invasion stage. Therefore, RBC membrane deformations are potentially induced by a passive mechanism, such as parasite adhesion.

In this paper, we focus on the passive compliance hypothesis [20] which assumes that RBC deformations and parasite alignment result from parasite adhesion interactions rather than from some active mechanism. Thus, our central question is whether parasite alignment can be explained purely by the passive compliance hypothesis. In contrast to the recent simulation study by [17], where RBC-parasite interactions are represented by a laterally smooth potential, the adhesion model presented here is based on discrete stochastic bonds between parasite and RBC membrane. This is a key step toward a realistic description of RBC-merozoite adhesion, since it eliminates the major shortcomings of the previous potential-based model such as unrealistically fast alignment times and the absence of dynamic membrane deformations. Even though receptor-ligand interactions which determine parasite alignment are largely not known, our bond-based interaction model still incorporates a few experimental details such as the range of adhesion forces and density of different agonists [5]. In particular, bonds of different lengths, i.e. long and short two-state bond interactions, are employed in the model. The bond-based parasite adhesion model generates a stochastic motion of the parasite at the RBC membrane, similar to that observed experimentally [16]. Furthermore, it results in alignment times which agree quantitatively with those measured in experiments [16, 21] and confirms the importance of membrane deformations for successful parasite alignment. The model is also used to investigate the effect of various bond properties, such as kinetic rates and bond density, on the parasite alignment process. Future investigations with this model can consider more realistic scenarios such as parasite adhesion and alignment under blood flow conditions.

The article is organized as follows. First, we introduce and calibrate our hydrodynamic model, where simulation parameters are tuned to quantitatively match several characteristics of the parasite motion at the RBC membrane from available experimental data by [16]. Then, RBC membrane deformations and alignment times are investigated for this reference parameter set and several cases of altered bond kinetics and densities. Finally, the effect of membrane stiffness on alignment times is studied.

## II. RESULTS

The RBC membrane is modeled as a network of *N*_rbc_vertices that are distributed uniformly on the membrane surface and connected by *N*_s_ springs [22–25]. Our RBC membrane model incorporates elastic and bending resistance, and its biconcave shape is obtained by constraining the total surface area and enclosed volume of the membrane.

Similar to the RBC, a parasite is modeled by *N*_para_ vertices distributed homogeneously on its surface. The egg-like shape of a merozoite (see Fig. 1(a)) is approximated as [6, 17]

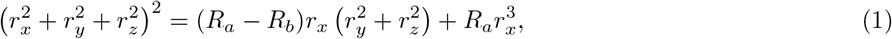

where *R_a_* = 1.5 μm and *R_b_* = 1.05 μm are diameters along the major and minor axes of the parasite, respectively. The parasite is much less deformable than the RBC, as no deformations of parasite body are visible in experiments [9, 16]. Therefore, the merozoite is considered to be a rigid body, whose dynamics can be described by equations involving force and torque on the parasite’s center of mass and directional vector [26].

**FIG. 1.**
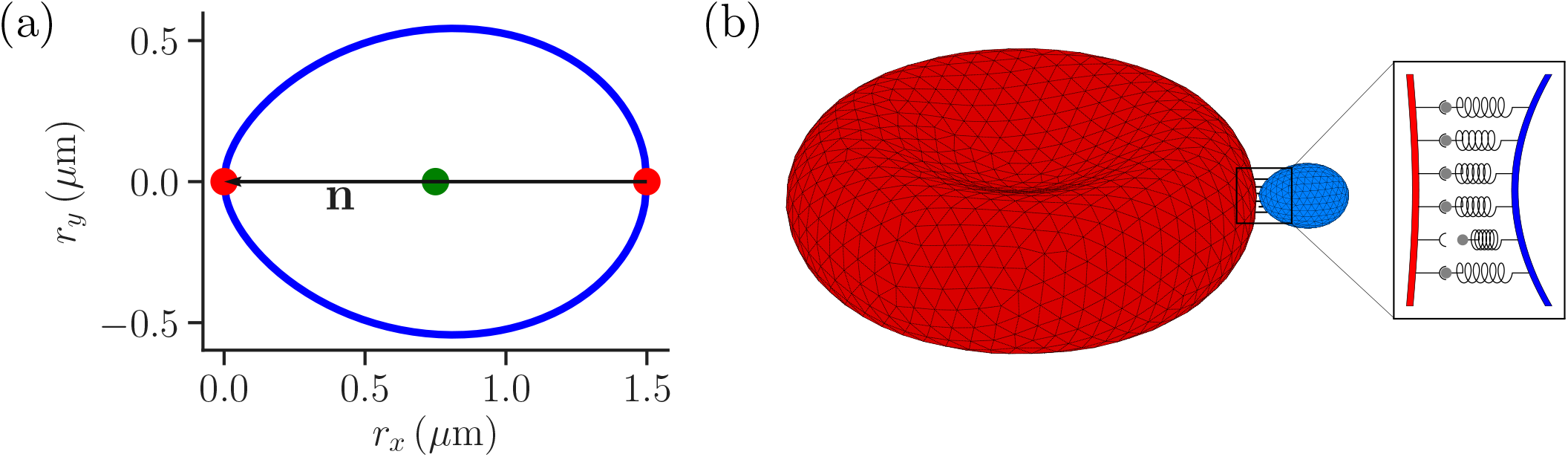
(a) Two-dimensional sketch of a parasite with a directional vector n from the parasite’s back at *r_x_* = 1.5 μm to its apex at *r_x_* = 0. (b) Three-dimensional triangulated surfaces of a RBC (red) and a parasite (blue). Bonds between the parasite and RBC can form within the contact zone which is illustrated by a magnified view, where discrete receptor-ligand interactions (or bonds) are sketched. A receptor-ligand bond can form with a constant on-rate *k*_on_ and break with a constant off-rate *k*_off_.

Both RBC and parasite are immersed in a fluid and the hydrodynamic interactions are modeled by the dissipative particle dynamics (DPD) method [27, 28]. The interaction of parasite and RBC membrane has two components. The first is an excluded-volume repulsion which is modeled by the repulsive part of the Lennard-Jones (LJ) potential with a minimum possible distance *σ*. The second represents adhesion which is modeled by discrete dynamic bonds between RBC and parasite vertices. A parasite vertex can form two different types of bonds: (i) long bonds with a maximum extension of 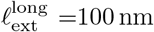 and (ii) short bonds with a maximum extension of 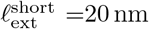, see *Methods* for more details.

To relate simulation units to physical units, a basic length scale is defined as the effective RBC diameter 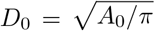 (*A*_0_ is the membrane area), an energy scale as *k*_B_*T*, and a time scale as RBC membrane relaxation time 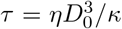, where *η* is the fluid viscosity and *κ* is the bending rigidity of the membrane. All simulation parameters in model and physical units are given in Tables I and II in *Methods* section. Average properties of a healthy RBC correspond to *D*_0_ ≃ 6.5 μm with *A*_0_ = 133.5 μm^2^ and *τ* ≈ 0.92s for *κ* = 3 × 10^−19^J and *η* = 1 mPas.

**TABLE I.**
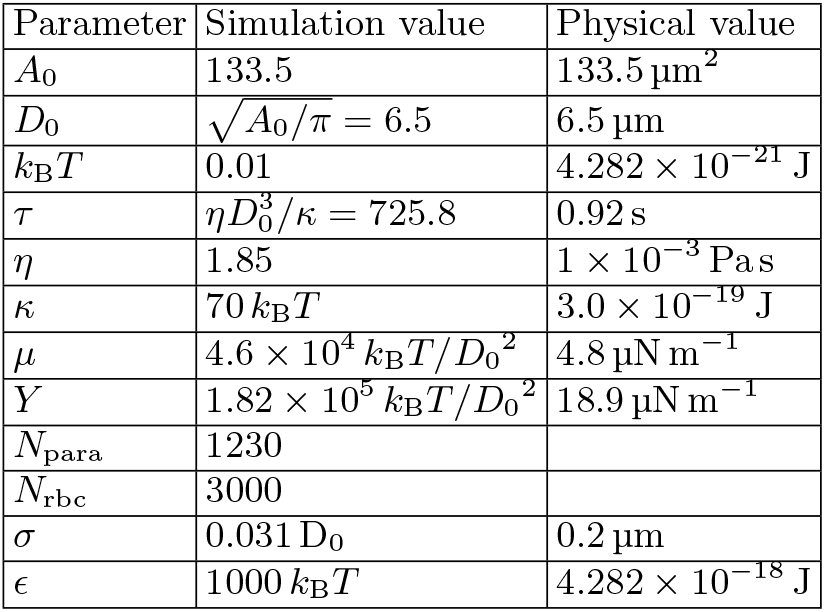
Simulation parameters given in both model and physical units. The effective RBC diameter 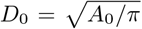 sets a basic length, the thermal energy *k*_B_*T* defines an energy scale, and RBC relaxation time 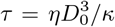 sets a time scale in the simulated system, where Ā0 is the RBC surface area, *κ* is the bending rigidity, and *η* is the fluid dynamic viscosity. The values of bending rigidity *κ*, shear *μ* and Young’s *Y* moduli are chosen such that they correspond to average properties of a healthy RBC. Parameters *σ* and *ϵ* correspond to RBC-parasite excluded-volume interactions represented by the purely repulsive LJ potential in Eq. (11).

**TABLE II.** List of bond parameters that are used to calibrate displacement of the parasite at the RBC membrane in simulations (see Movie S1) against available experimental data [16], as shown in Fig. 2(c). The parameter values in simulations are given in terms of the length scale *D*_0_, energy scale *k*_B_*T*, and timescale 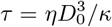. The densities of long and short bonds are given in terms of parasite vertex density *ρ*_para_ ≃ 270μm^−2^. Note that *ρ*_long_ + *ρ*_short_ = *ρ*_para_ in all simulations.

| Parameter | Simulation value | Physical value |
| --- | --- | --- |
| $\ell_{\text{ext}}^{\text{long}}$ | $0.0154 D_0$ | 100 nm |
| $\ell_{\text{ext}}^{\text{short}}$ | $0.0031 D_0$ | 20 nm |
| $\rho_{\text{long}}$ | $0.4 \rho_{\text{para}}$ | $107 \mu\text{m}^{-2}$ |
| $\rho_{\text{short}}$ | $0.6 \rho_{\text{para}}$ | $161 \mu\text{m}^{-2}$ |
| $k_{\text{on}}^{\text{long}}$ | $36.3 \tau^{-1}$ | $39.6 \text{ s}^{-1}$ |
| $k_{\text{on}}^{\text{short}}$ | $290.3 \tau^{-1}$ | $317.0 \text{ s}^{-1}$ |
| $k_{\text{off}}$ | $72.58 \tau^{-1}$ | $79.2 \text{ s}^{-1}$ |
| $\lambda_{\text{long}}$ | $2.46 \times 10^4 k_B T/D_0^2$ | $2.57 \mu\text{N m}^{-1}$ |
| $\lambda_{\text{short}}$ | $0.82 \times 10^4 k_B T/D_0^2$ | $0.856 \mu\text{N m}^{-1}$ |

To better understand the effect of various adhesion properties on parasite alignment, several parameters such as bond formation and rupture rates and relative bond densities are varied. For each fixed parameter set, a number of simulations are performed and the results are combined and/or averaged, which is necessary due to the stochastic nature of bond-based interaction as well as thermal fluctuation effects within the fluid. Note that each simulation is performed for a different random choice of membrane vertices which form long and short bonds, while their relative densities remain fixed, see *Methods*.

### A. Calibration of RBC-parasite interactions

A parasite adhered to the RBC membrane generally exhibits a stochastic (or diffusive-like) motion observed experimentally [16], which is controlled by the receptor densities *ρ*_long_ and *ρ*_short_ and the bond formation 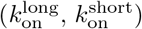 and rupture (*k*_off_) rates that are currently not known. Nevertheless, available experiments [5] suggest that the number of short bonds in RBC-merozoite interaction is lager than the number of long bonds, which is reflected in the receptor densities *ρ*_long_ and *ρ*_short_ assumed for our parasite model (see Table II). To calibrate RBC-parasite interactions, the stochastic dynamics of the parasite adhered to the RBC membrane (see Movie S1) is quantified by its fixed-time displacement, which is measured by tracking the distance Δ*d* traveled by the parasite at fixed intervals of time Δ*t*, see Fig. 2(a). Particle tracking is employed to measure Δ*d* from available experiments [16], where Δ*t* is selected to be 1 s, which is the time resolution of the experimental videos. Only time ranges, within which parasites remain visible and the RBC is not moving much, are included in the analysis.

**FIG. 2.**
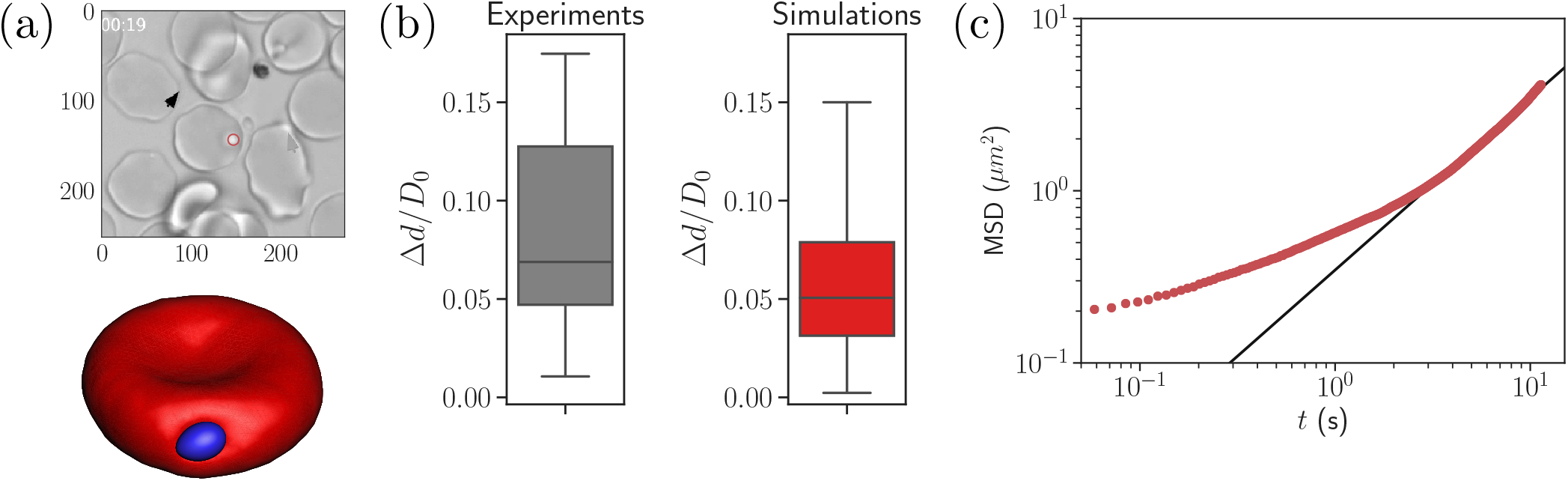
(a) A time instance of parasite motion at RBC membrane from an experimental movie [16] (top) and simulation (bottom), see also Movie S1. To obtain the distribution of merozoite fixed-time displacements, the marked parasite (red circle) is tracked over the course of its interaction with the RBC membrane. (b) Comparison between experimental and simulated fixed-time displacements (Δ*d*) of the parasite at RBC membrane, which is normalized by the effective RBC diameter 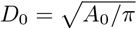 calculated from the membrane area *A*_0_. By adapting the interaction parameters, the displacement distribution in simulations is calibrated against the experimental distribution. The resulting interaction parameters for our model can be found in Table II. (c) Mean squared displacement (MSD) of a parasite from simulations as a function of time. The black solid line marks a diffusive regime with MSD ~*t*. Note the subdiffusive dynamics for short times, less than about 1 s.

Figure 2(b) compares experimental and simulated characteristics of fixed-time displacements for the interaction parameters given in Table II. This set of parameters is obtained by varying *ρ*_long_, *ρ*_short_, 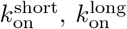, and *k*_off_ until a good agreement between experimental and simulated parasite displacements is reached. However, the maximum extensions of long and short bonds remain fixed at 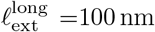 and 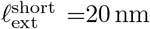 in this calibration procedure. The variance of experimental displacements in Fig. 2(b) is larger than that in simulations due to a limited sample size of experimental data.

To further characterize the parasite motion on the RBC membrane, the mean-squared displacement (MSD) of the parasite’s center of mass is computed in simulations and shown in Fig. 2(c). At long enough times *t* ≳ 3 s, the parasite exhibits diffusive-like motion, indicated by a linear increase of the MSD curve with time. For shorter timescales, the MSD of parasite motion shows a transient anomalous subdiffusion, which may occur, for instance, in the case of sticky particle dynamics with alterations between sticking (i.e., stopping its motion for some time) and diffusing states [29, 30]. The transient sticky dynamics is an appropriate description for an adhered parasite, where sticking periods correspond to time intervals within which no bonds are formed or ruptured. The diffusive-like dynamics is governed by the number of bonds *n_b_* and their on- and off-rates, as an adhered particle becomes slower and eventually gets arrested when *n_b_* is increased and the rates are decreased [31].

### B. Parasite alignment

Recent experiments suggest that a successful RBC invasion strongly correlates not only with the distance between parasite apex and RBC membrane, but also with a perpendicular alignment of the merozoite toward the cell membrane [7]. Furthermore, the junctional (invasion initiating) interaction range *r*_junc_ of the parasite’s apex is known to be around 10 nm [5]. Based on these observations, we define two quantities, (i) the apex distance *d*_apex_ from the RBC membrane, and (ii) the alignment angle *θ* that characterizes parasite orientation, both sketched in Fig. 3(a). Here, *d*_apex_ is defined as the distance between the parasite apex and the nearest membrane vertex,

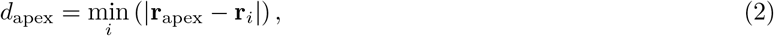

the alignment angle *θ* as the angle between the parasite’s directional vector **n** and the normal **n**^face^ of a triangular face whose center is closest to the apex,

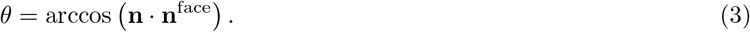

**FIG. 3.**
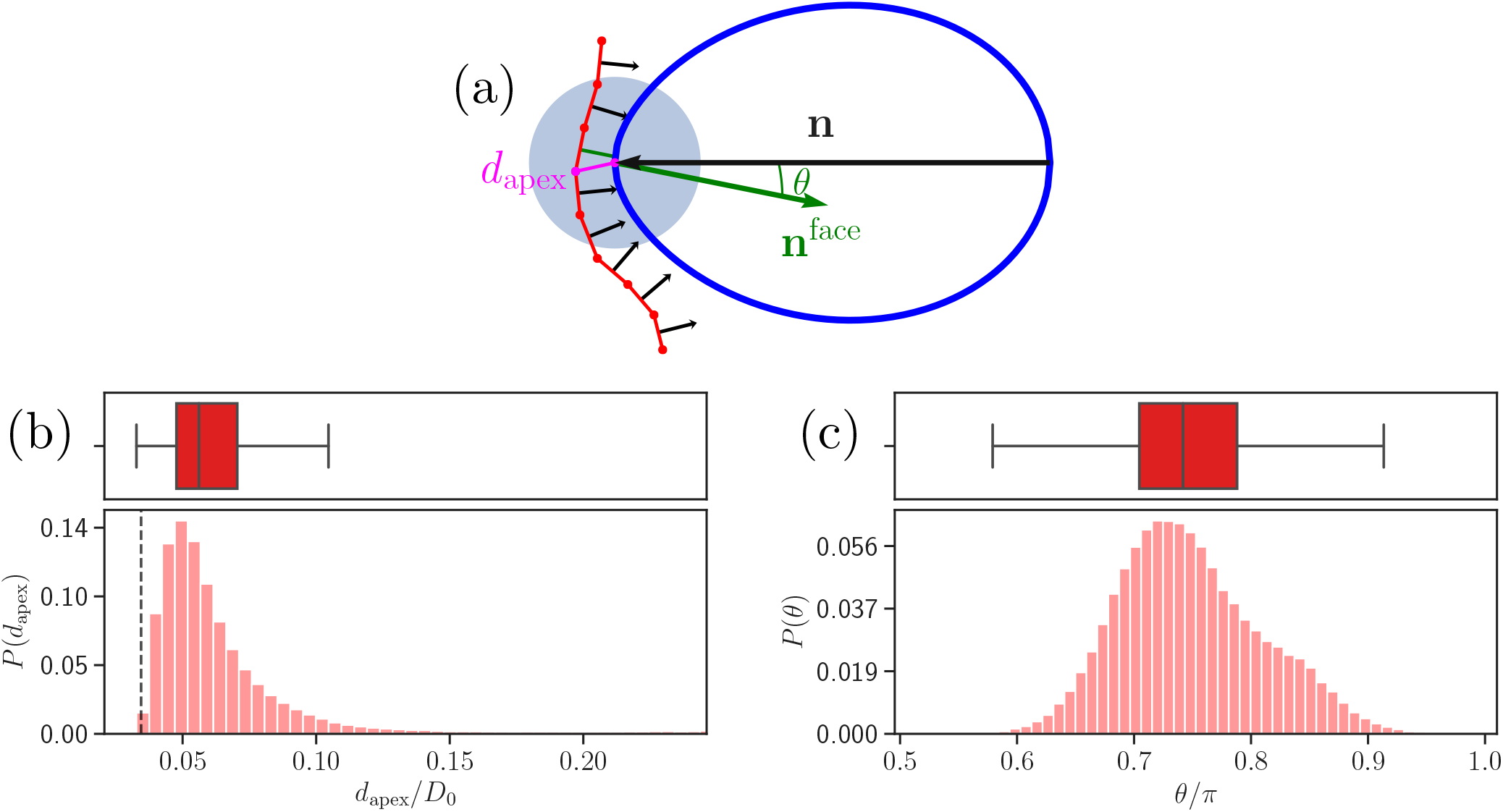
(a) Sketch of apex distance *d*_apex_ and alignment angle *θ*. *d*_apex_ is defined as a distance (magenta line) between the parasite’s apex and the closest vertex of RBC membrane. The alignment angle *θ* corresponds to the angle between the parasite’s directional vector (black arrow) and the normal vector **n**^face^(green arrow) of a triangular face whose center is closest to the apex. (b) & (c) Probability distributions of the apex distance *d*_apex_/*D*_0_ and the alignment angle *θ/π*. Data are obtained for parameters shown in Table II. The dashed line in the apex distance distribution indicates the minimum possible distance *σ* of the repulsive LJ potential. Note that a good parasite alignment requires small values of *d*_apex_/*D*_0_ and values of *θ/π* close to unity.

Note that the search for the face closest to the apex is performed only within a cutoff range for long bonds (i.e. within 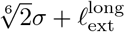) from the apex, indicated by the blue circle in Fig. 3(a).

Figure 3(b,c) shows distributions of apex distance *d*_apex_ and alignment angle *θ* for the calibrated RBC-parasite interactions. Both characteristics are represented by distributions as the merozoite moves stochastically at the membrane surface. Minimum values of *d*_apex_ in Fig. 3(b) correspond to the parasite’s apex being very close to the membrane (i.e., *d*_apex_ ≈ *σ*), whereas maximum values generally represent a configuration where the parasite is adhered sideways to the RBC. Furthermore, low values of *θ* in Fig. 3(c) characterize the sideways adhesion orientation, while large values of *θ* represent a good alignment configuration. Note that an ideal merozoite alignment would be achieved if *d*_apex_ is less than *σ* + *r*_junc_ (*r*_junc_ = 10nm) and the alignment angle is *θ* ≈ *π*. Due to a discrete representation of the membrane, perfect alignment is unlikely, which requires to slightly relax these conditions. Therefore, we define a successful parasite alignment by the criteria

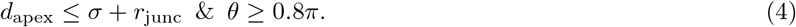

In experiments, merozoite alignment times are measured as time intervals between initial parasite adhesion and the beginning of invasion [16]. Similarly, alignment time in simulations is calculated as the time required for the parasite to meet the alignment criteria in Eq. (4) starting from an initial adhesion contact. Figure 4(b) presents a distribution of alignment times from 86 statistically independent DPD simulations for the reference RBC-parasite interactions in Table II. The alignment times range between 1 s and 26 s with an average value of 9.53 s. For comparison, the average alignment time was reported to be 16 s by [16], and the range of alignment times between 7 s and 44 s was found by [21], which agree reasonably well with our model predictions.

**FIG. 4.**
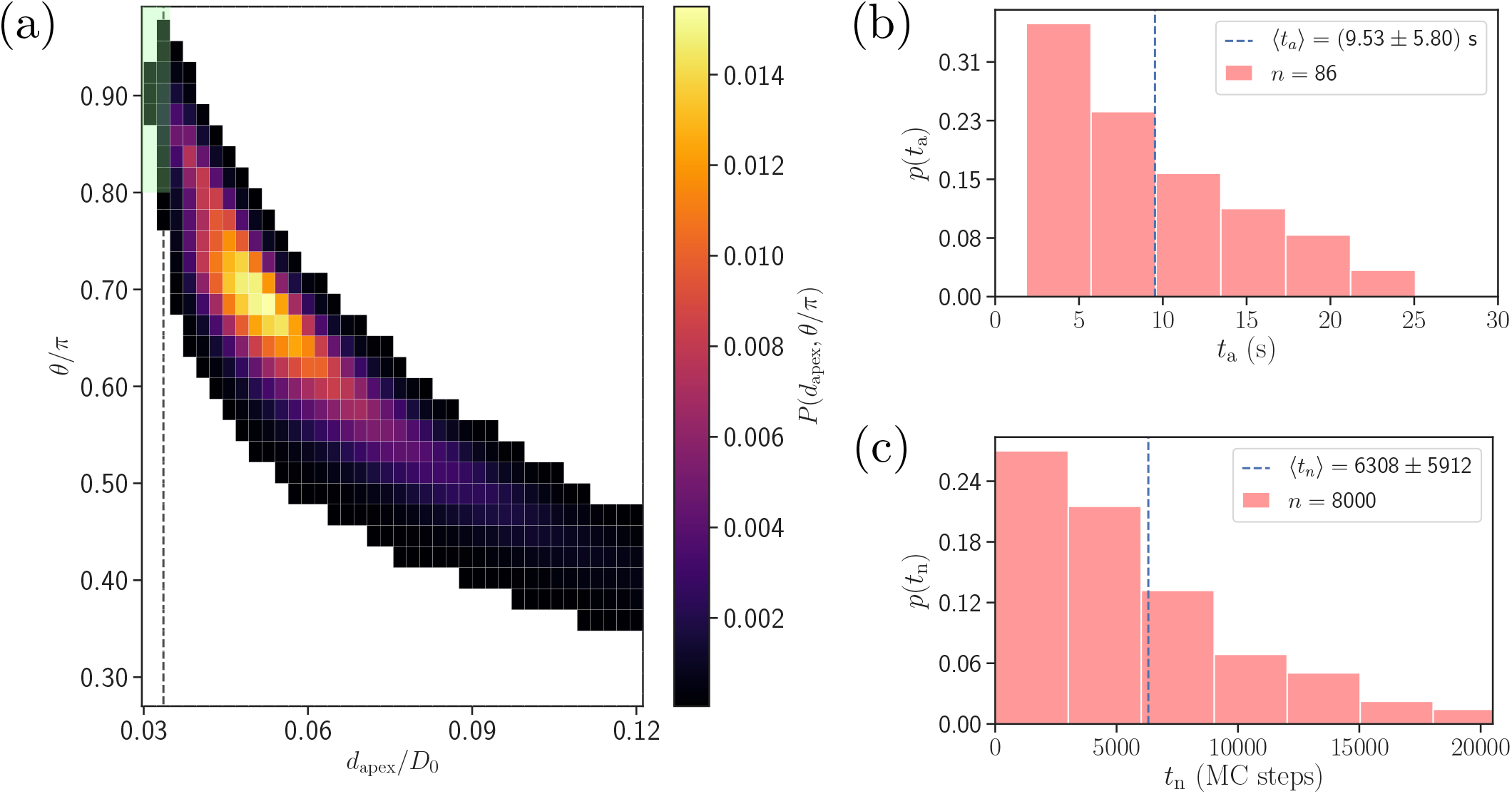
(a) Two-dimensional probability map as a function of *d*_apex_ and *θ*. Each bin represents a single alignment state and the color corresponds to probability of that state. The dark green area represents the criteria for a successful alignment. (b) Distribution of alignment times *t*_a_ obtained from 86 statistically independent DPD simulations. *t*_a_ is defined as a time interval between the start of a simulation and the instance when the alignment criteria for dapex and *θ* in Eq. (4) are met. The average alignment time is equal to 〈*t*_a_〉 ≃ 9.53 s. (c) Alignment time distribution from MC sampling using the probability map in Fig. 4(a). The alignment time is defined as a number of MC steps needed to satisfy the alignment criteria, as the MC procedure does not have an inherit timescale. Note that the sample size in MC modeling is much larger than that in Fig. 4(b).

Note that the sample size (about 100) in simulations is limited by the computational cost. A single simulation, corresponding to a total physical time of about 26s, requires approximately 168 core hours on the supercomputer JURECA [32] at Forschungszentrum Jϼlich. Therefore, a direct brute-force approach for the investigation of the effect of various parameters on the parasite alignment time is not feasible. To overcome this problem, Monte-Carlo (MC) sampling (see *Methods* for details), which is based on a two-dimensional probability map of parasite alignment characteristics (*d*_apex_, *θ*) illustrated in Fig. 4(a), is employed to determine the differences in alignment times for various parameter sets. Such a probability map is computed from several direct DPD simulations of RBC-parasite adhesive interactions. Then, the MC procedure is used to model stochastic jumps between neighboring alignment states (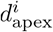, *θ^j^*) within the probability map, starting from a randomly selected initial state and continuing until the alignment criteria in Eq. (4) are met, and the number of MC steps represents the alignment time. Distribution of alignment times *t*_n_from the MC sampling is shown in Fig. 4(c) for the reference parameter set. Clearly, the distributions obtained by direct (Fig. 4(b)) and MC (Fig. 4(c)) simulations are very similar, verifying the reliability of the MC approach. Note that alignment times t**n** from MC sampling are measured in terms of MC steps, since MC simulations do not have an intrinsic timescale. The average alignment time for the reference parameter set is denoted as 〈*t*_*n*,ref_〉 and assumed to be equivalent to 9.53 s, the average alignment time from direct DPD simulations of RBC-parasite adhesion. This implies that 10^4^ MC steps correspond to about 15 s.

### C. Membrane deformation and parasite dynamics

A recent simulation study by [17] with a laterally homogeneous adhesion potential has demonstrated that the deformation of RBC membrane is crucial for a successful parasite alignment. Here, we show that bond density and kinetics not only control the parasite motion at the membrane surface, but also directly affect membrane deformation. To quantify the strength of membrane deformations, a change in total energy between the deformed state and the equilibrium state of the RBC membrane is computed as [17]

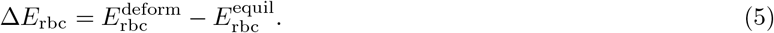

Figure 5 shows temporal changes in deformation energy, number of bonds, head distance, and alignment angle for the reference case. Two major contributions to the deformation energy (i.e. elastic stretching Δ*E*_sp_ and bending Δ*E*_bend_ energies) indicate that membrane deformation is very dynamic and has a strong variability in its intensity. This is due to the dynamic formation and dissociation of long and short bonds between the merozoite and RBC membrane. An interesting observation is that the head distance and alignment angle in Fig. 5 fluctuate around some average values, indicating that the parasite has a preferred orientation, which is consistent with a peak in the probability map in Fig. 4(a). In fact, the average values of *d*_apex_ and *θ* correspond to a parasite orientation that has a maximum contact area with the membrane, and are determined by the parasite’s egg-like shape. Furthermore, the fluctuations of *d*_apex_ and *θ* from their average values represent parasite motion toward its apex or bottom due to stochastic bond dynamics. Thus, the parasite dynamics at the membrane can be described as a superposition of the rolling motion around its directional vector with a preferred orientation and intermediate fluctuations of parasite orientation toward its apex or the bottom. The rotational motion around the directional vector is preferred because it is not associated with a significant energy cost, while fluctuations in the orientation toward the merozoite’s apex or bottom have an energy penalty.

**FIG. 5.**
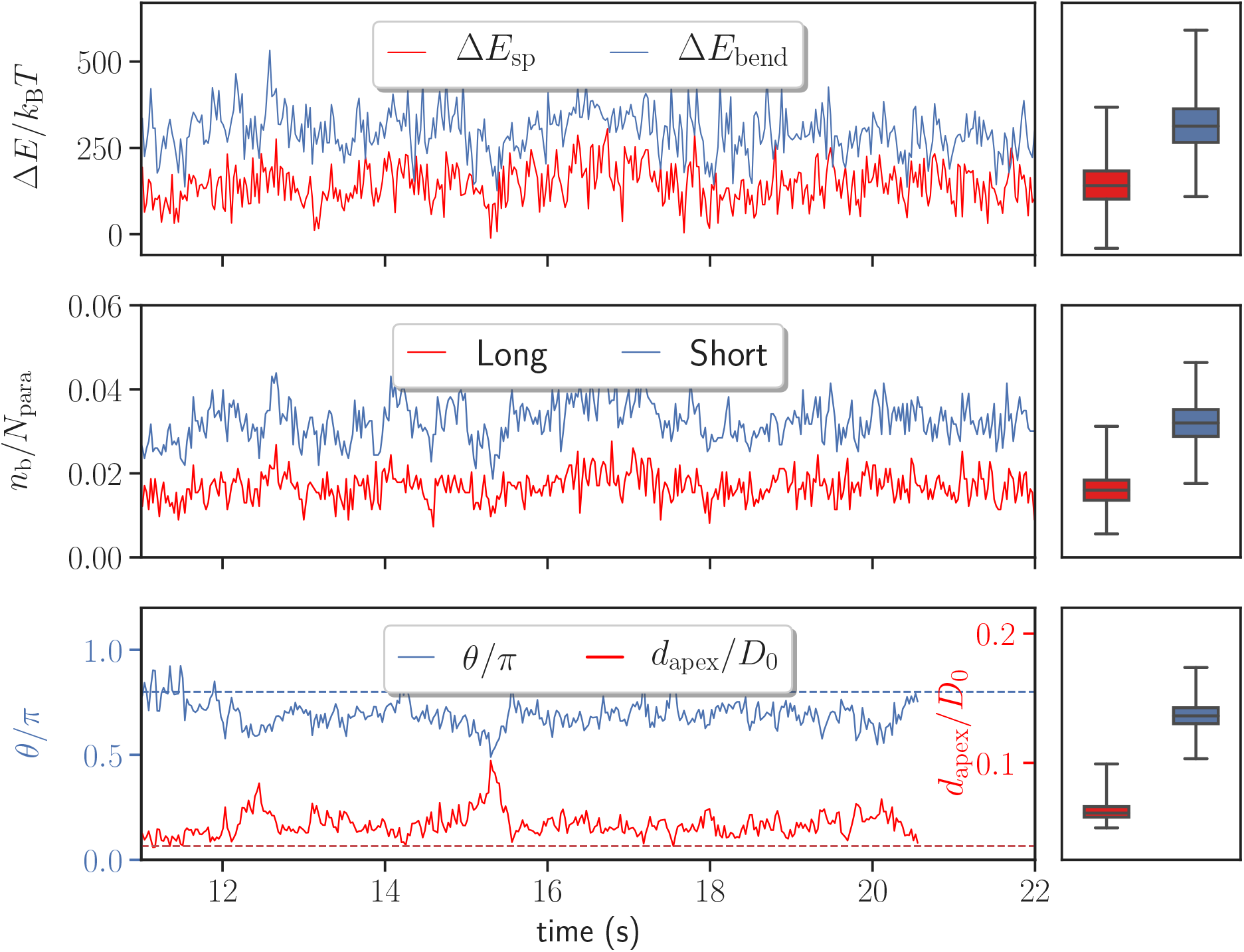
Variations in stretching Δ*E*_sp_ and bending Δ*E*_bend_ energies, the number of bonds *n_b_*, the head distance *d*_apex_, and the alignment angle *θ* as a function of time for the default parameter set given in Table II. Temporal changes in the number of bonds are shown for both long and short bond types. The dashed lines in the bottom plot correspond to the alignment criteria in Eq. 4. For all quantities, the corresponding averages and variances represented by box plots are depicted on the right.

The dynamic adhesive behavior of the parasite in the current stochastic model is in striking contrast to the previous adhesion model [17] based on a homogeneous interaction potential between the two cells, where no dynamic deformations were observed. A qualitative correspondence between these two models can be understood by considering a ratio *k*_on_/*k*_off_ = exp (Δ*U_b_*/*k*_B_*T*), where Δ*U_b_* is the binding energy of a single bond [33, 34]. Thus, the ratio *k*_on_/*k*_off_ directly controls the average number of bonds 〈*n_b_*〉 and the strength of adhesion, which are correlated with RBC deformation energy Δ*E*_rbc_. Similarly, in the parasite adhesion model with a homogeneous interaction potential [17], the strength of adhesion potential controls membrane deformations. Even though average membrane deformations can be compared for these two models, the stochastic bond-based adhesion model results in a very different diffusive-like dynamics of the parasite, which is governed by *n_b_* and the off-rate *k*_off_ [31]. A significant increase of *n_b_* and/or a decrease of *k*_off_ would lead to parasite arrest, which can be compared well with the model based on a homogeneous interaction potential [17].

There exist three different timescales which might be relevant for the parasite alignment: (i) bond lifetime *τ_b_* ≃ 1/*k*_off_, (ii) membrane deformation time on the scale of parasite size 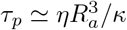, and (iii) rotational diffusion time of the parasite 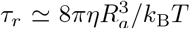. These characteristic times are *τ_b_* ≈0.013s, *τ_p_* ≈0.011s, and *τ_r_* ≈20s computed from the model parameters given in Tables I and II in Methods section. There is a clear separation of timescales between *τ_r_* and both *τ_b_* and *τ_p_*, indicating that the rotational diffusion of the parasite is too slow to have a significant effect on merozoite alignment. Furthermore, *τ_b_* and *τ_p_* are comparable in magnitude, suggesting that both bond dynamics and membrane deformations are important for the alignment process. It is also interesting to note that the ratio *τ_p_/τ_r_ = k*_B_*T*/(8*πκ*) ≈ 6× 10^−4^ depends only on the bending rigidity *κ*. This means that membrane deformation will always represent a dominating timescale over the rotational diffusion of the parasite, independently of the parasite size and the viscosity of suspending medium.

### D. Effect of bond properties on parasite alignment

To better understand the dependence of merozoite alignment on bond kinetics, the off-rate *k*_off_ is varied for two ratios 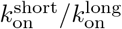 of short and long bond on-rates. Figure 6 presents the parasite’s fixed-time displacement, deformation energy, and average alignment times as a function of 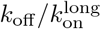. A lower ratio of 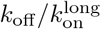 (i.e. a lower *k*_off_) leads to stronger adhesion and thereby stronger membrane deformations (see Fig. 6(b) and Movie S2), consistently with the discussion above. For small 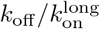 values, membrane deformation energies can reach up to 2000 *k*_B_*T*, whereas large values of *k*_off_ result in Δ*E*_rbc_ ≈ 100 *k*_B_*T*. The main reason is that low values of *k*_off_ lead to a significant increase in the lifetime of individual bonds, allowing the parasite to form more bonds and thereby increase its adhesion energy and induce larger membrane deformations. Similarly, large values of *k*_off_ decrease the bond lifetime, resulting in a decrease in the adhesion energy. For instance, in case of 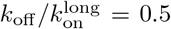, the parasite forms on average about 200 bonds, whereas for 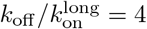, the average number of bonds is approximately 15. Furthermore, a larger on-rate for the short bonds yields a slight increase in the strength of membrane deformations in comparison to a smaller 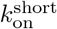.

**FIG. 6.**
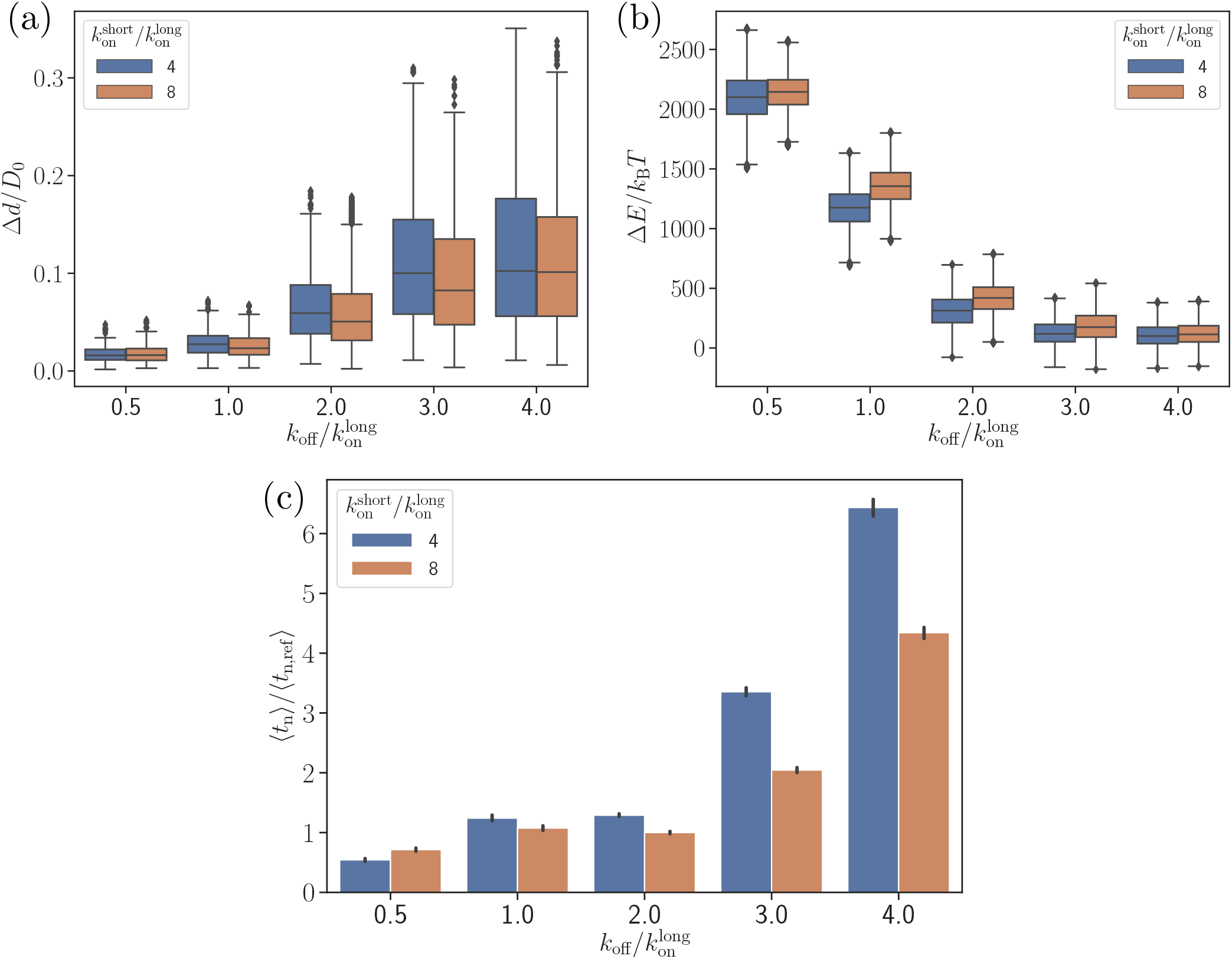
Effect of the off-rate *k*_off_ on (a) the parasite’s fixed-time displacement, (b) RBC deformation energy, and (c) alignment time. Since the off-rate controls the lifetime of bonds, a smaller off-rate results in a stronger adhesion, a lower parasite displacement, and a faster alignment time.

Figure 6(b,c) shows that there is a clear correlation between the level of membrane deformations and average alignment time. For example, for off-rates 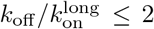, the alignment times are comparable with those for the reference parameter case, while for off-rates 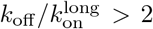, there is a strong increase in alignment times, which is correlated with insignificant membrane deformations. A shorter alignment time for 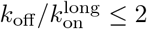 is due to the partial wrapping of the parasite by the RBC membrane, which is consistent with the previous study by [17] that demonstrates the importance of membrane deformation for merozoite alignment. Note that the fixed-time displacement Δ*d* in Fig. 6(a) significantly increases with *k*_off_ due to a weaker adhesion. This seems to imply that the parasite alignment proceeds faster for 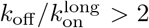. However, as it is evident from Fig. 6(c), this simple expectation is not applicable here, indicating that a faster motion of the parasite at the RBC surface may not result in a faster alignment.

Another parameter, which may significantly affect parasite alignment, is the density of long bonds *ρ*_long_. For the reference parameter set, *ρ*_long_ is chosen to be *ρ*_long_/*ρ*_para_ = 0.4, so that *ρ*_short_/*ρ*_para_ = 0.6. Figure 7 presents the number of short and long bonds as well as parasite alignment times as a function of *ρ*_long_/*ρ*_para_. For the density *ρ*_long_/*ρ*_para_ = 0.1, the value of 〈*t_n_*〉 is omitted, as the alignment criteria in Eq. (4) have not successfully been met during the entire course of direct simulations, yielding the probability of parasite alignment in MC sampling to be zero. For densities *ρ*_long_/*ρ*_para_ ≥ 0.3, both bond numbers and alignment times remain nearly independent of *ρ*_long_. However, the average alignment time for *ρ*_long_/*ρ*_para_ = 0.2 is about 30 s which is roughly three times longer than for the reference case. Note that 30 s is longer than the total length (≈26 s) of direct simulations. Nevertheless, parasite alignment has occurred in some of these simulations, resulting in a small non-zero probability of merozoite alignment and a relatively long 〈*t_n_*〉 calculated through the MC sampling. The fact that 〈*t_n_*〉 for *ρ*_long_/*ρ*_para_ = 0.2 is longer than the total time of direct simulations means that the probability of parasite alignment is likely overestimated, indicating that the average alignment time should be even longer than 30 s. An increase of with decreasing values of *ρ*_long_ is consistent with a significant decrease in membrane deformations. For off-rates *k*_off_ < 72.6 *τ*^−1^, the trends illustrated in Fig. 7 remain qualitatively the same.

**FIG. 7.**
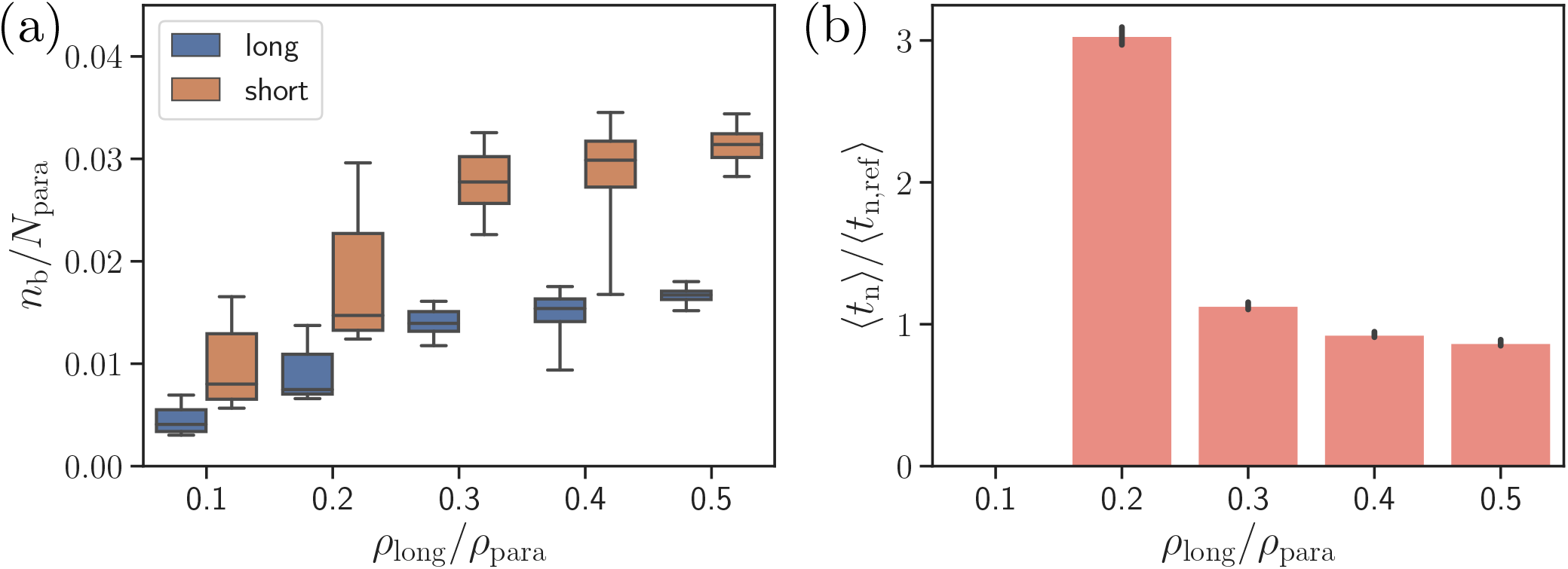
Effect of the density of long bonds *ρ*_long_ on parasite alignment. (a) Number of short and long bonds and (b) parasite alignment times as a function of *ρ*_long_/*ρ*_para_. Note that *ρ*_long_ + *ρ*_short_ = *ρ*_para_ remains constant in all simulations. Here, the bond kinetic rates are 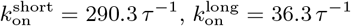, and *k*_off_ = 72.6 *τ*^−1^. In case of *ρ*_long_/*ρ*_para_ = 0.1, parasite alignment time could not be computed through the MC sampling, since merozoite alignment has never occurred in direct simulations.

### E. Effect of RBC rigidity

To investigate the effect of RBC rigidity on the alignment of a merozoite, we consider a nearly rigid cell membrane by increasing both bending rigidity and Young’s modulus by two orders of magnitude in comparison to a healthy RBC. Such a rigid RBC shows no significant membrane deformations for the reference interaction parameters given in Table II, see Movie S3. Comparison of parasite fixed-time displacements and alignment times for flexible and rigid membranes is shown in Fig. 8 for two different values of *k*_off_. Clearly, larger RBC rigidity leads to much longer parasite alignment times (see Fig. 8(b)), emphasizing again the importance of membrane deformations for merozoite alignment. For off-rates 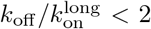, parasite alignment at the surface of a rigid RBC is not achieved within the course of the simulation. As the off-rate increases, alignment time at the rigid membrane becomes comparable with that for the flexible membrane, because large enough *k*_off_ values do not result in strong membrane deformations even for the flexible RBC. Thus, for large off-rates, the parasite’s alignment solely relies on its rotational diffusion controlled by the kinetic bond rates.

**FIG. 8.**
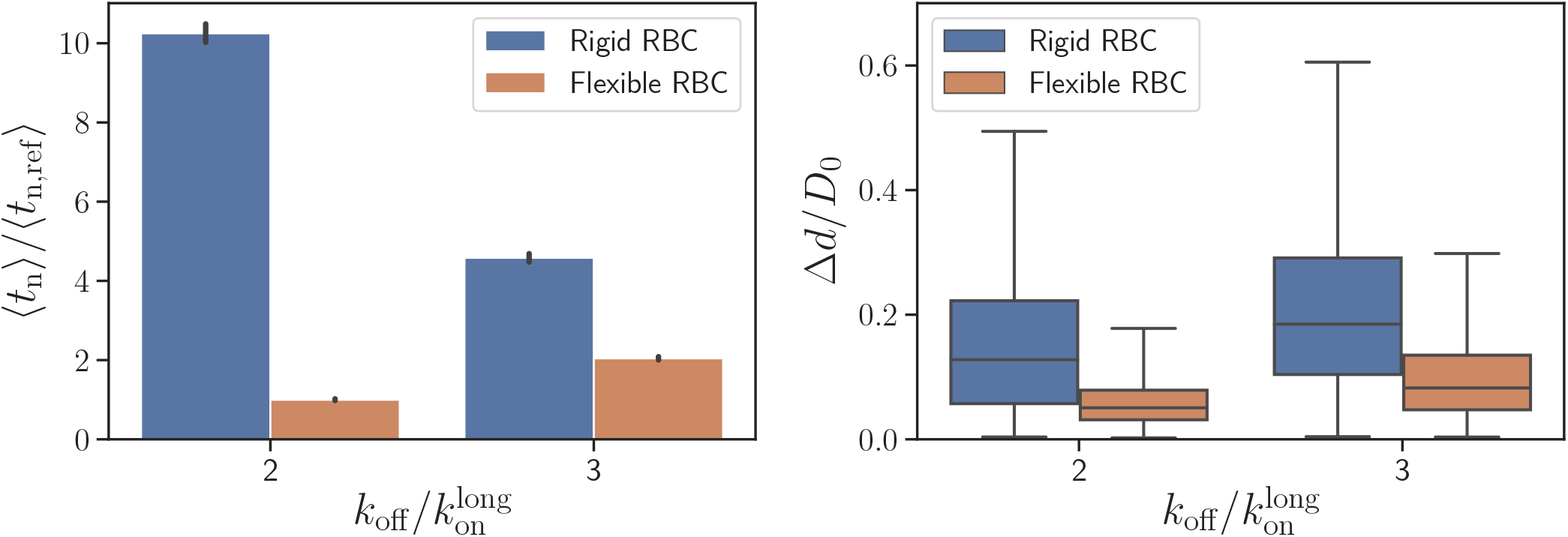
Effect of RBC membrane rigidity on (a) alignment time and (b) parasite fixed-time displacement for different off-rates *k*_off_. Note that for a rigid RBC with 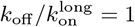, parasite alignment time could not be computed through the MC sampling, as the alignment criteria have never been met in direct simulations.

Figure 8(a) presents a comparison of parasite fixed-time displacements at the flexible and rigid membranes. In both cases, parasite displacements increase with increasing *k*_off_, as expected. However, the displacement at the rigid membrane is larger than at the flexible membrane (for visual comparison, see Movies S1 and S3), because the merozoite forms less bonds at the rigid surface. For the same reason, the variance of parasite displacements is larger for the rigid RBC than for the flexible RBC. Note that an increase in *k*_off_ results in an increase of fixed-time displacement and a decrease of alignment time for the rigid membrane, whereas for flexible RBC, an increase in off-rate leads to an elevation of both fixed-time displacement and alignment time. This implies that for a rigid RBC, fast kinetics or weak adhesion are favorable for a quick alignment. In contrast, for a flexible RBC, slow kinetics or strong adhesion are advantageous for fast alignment, since the parasite employs RBC deformation for efficient alignment by partial membrane wrapping.

## III. DISCUSSION AND CONCLUSIONS

We have investigated the alignment of a merozoite at RBC membrane using a realistic two-state bond-dynamics model for parasite adhesion. Motivated by experiments [5], parasite adhesion is modeled by two bond types, with long and short extension. Since RBC-parasite interactions and the corresponding bond kinetics are experimentally not yet well characterized, the calibration of bond parameters is based on parasite fixed-time displacement at the membrane from existing experiments [16], which is in the range of 0.3 – 0.8 *μ*m. The presented model is able to reproduce quantitatively experimentally measured alignment times. Simulated alignment times are in the range between a few seconds and 26s, while the analysis of experimental videos by [16] yields an average alignment time of 16 s. Another independent experimental study by [21] reports alignment times in the range between 7 and 44 s, which agree relatively well with our simulation predictions. In addition to the good agreement between simulated and experimental alignment times, our model reproduces well dynamic RBC membrane deformations frequently observed in experiments [12, 13, 15].

Our main result is that parasite alignment is mediated by RBC membrane deformations and a diffusive-like dynamics due to the stochastic nature of parasite-membrane interactions. Average number of bonds 〈*n_b_*〉 between the parasite and the membrane is governed by the ratio *k*_on_/*k*_off_ = exp (Δ*U_b_*/*k*_B_*T*) that is connected to the binding energy Δ*U_b_* of a single bond and determines the strength of membrane deformations. Our results show that membrane deformations speed up the alignment through partial wrapping of the parasite, facilitating a contact between the parasite apex and the membrane. This conclusion is consistent with the previous simulation study [17], where merozoite adhesion has been modeled by a laterally homogeneous interaction potential whose strength controls RBC deformations. The importance of membrane deformation is also corroborated by simulations of parasite alignment at a rigid RBC, which show a drastic increase in alignment times. For a rigid membrane, the parasite alignment depends mainly on bond lifetime (i.e., *τ_b_* ≃ 1/*k*_off_), indicating that a low *k*_off_ or large bond lifetime may significantly decelerate the parasite’s rotational motion, and hence, increase its alignment time drastically. This conclusion agrees well with a recent simulation study [31] on the dynamics of two adhered colloids, whose effective rotational diffusion is governed not only by 〈*n_b_*〉 but also by *τ_b_*. Clearly, *τ_b_* is also important for parasite dynamics at a deformable RBC, in addition to the membrane relaxation time on the scale of parasite size *τ_p_*. The poor alignment of the merozoite at a stiff membrane can be a contributing factor, limiting parasite invasion. For example, infected RBCs in malaria become significantly stiffer than healthy cells [35, 36], limiting secondary invasion events. Furthermore, an increased RBC membrane stiffness is relevant in many other diseases, such as sickle cell anemia [37], thalassemia [38], and stomatocytosis [39], whose carriers are generally less susceptible to malaria infection.

For large values of *k*_off_, the parasite is not able to induce strong deformations even at a flexible membrane, so that the alignment times at rigid and deformable RBCs become comparable, and the alignment is governed solely by a diffusive-like rotational dynamics. The diffusive-like motion of the parasite at the membrane surface is facilitated by stochastic formation/dissociation of bonds between the two cell surfaces, and leads occasionally to a successful alignment. Therefore, our model is also able to explain the possibility of RBC invasion by a merozoite without preceding membrane deformations, which is observed much less frequently than the invasion preceded by significant RBC deformations [16]. Note that the RBC-parasite adhesion model based on a laterally homogeneous interaction potential [17] predicts the complete failure of parasite alignment without significant membrane deformations, because it does not capture a diffusive-like rotational dynamics of the parasite. Thus, the bond-based model is more appropriate for the representation of RBC-parasite interactions.

Even though the bond parameters in Table II were calibrated by the parasite fixed-time displacement obtained from experiments [16], such choice is likely not unique as some other set of parameters (e.g., receptor/ligand densities, kinetic rates) may lead to statistically similar displacement characteristics. Nevertheless, it is important to emphasize that the discrete bonds in simulations should be thought of as “effective” bonds, which likely represent a small cluster of real molecular bindings. Furthermore, since the parasite displacement is mainly controlled by the bond kinetics, this calibration procedure is rather robust in identifying an appropriate range of bond properties. Another important aspect of this model is the necessity of sufficiently long bonds to facilitate stochastic motion of the parasite at RBC surface. Simulations with only short bonds show that the parasite is quickly arrested, which is similar to the model for RBC-parasite adhesion based on a laterally homogeneous interaction potential [17]. Therefore, the long bonds serve as leverages for parasite motion at the membrane. Electron microscopy images of adhered parasites [5] suggest that the density of long bonds can be as low as 5 – 10%. However, the density of long bonds in our simulations is limited by the resolution of both the RBC and parasite to be larger than about 20%. A much finer membrane model would alleviate this limitation, but it would be prohibitively expensive computationally. Note that such heterogeneous systems of receptors exist in other biological systems as well. For example, during leukocyte binding in the microvasculature, both selectin and integrin molecules participate in adhesion and work synergistically, even though they have distinct functions [40]. Furthermore, infected RBCs in malaria adhere to endothelial cells via two distinct ligands, ICAM-1 and CD-36, where binding with ICAM-1 exhibits a catch-like bond, while the interaction with CD-36 is a slip-like bond [41].

Several studies [3, 6, 42] about RBC-parasite interactions hypothesize the existence of an adhesion gradient along the parasite body, which is expected to facilitate alignment. Based on the RBC-parasite adhesion model with a laterally homogeneous interaction potential [17], it was shown that an adhesion gradient, where the potential strength increases toward the apex of a merozoite, generally accelerates parasite alignment. No definite conclusions about possible gradients can be made in the context of that model, because even in the case of no adhesion gradients, it predicts very short alignment times of about two orders of magnitude smaller than measured experimentally. An introduction of adhesion gradients in our bond-based interaction model leads qualitatively to the following conclusions: (i) Weak adhesion gradients do not significantly disturb the stochastic motion of a parasite at RBC membrane, and have a negligible effect on the alignment. (ii) Strong adhesion gradients often result in a controlled direct re-orientation of the parasite toward its apex, suppressing the stochastic motion observed experimentally. These preliminary results do not permit a definite conclusion about the possible existence of adhesion gradients, as moderate adhesion gradients may exist and aid partially in the alignment process. Nevertheless, our model shows that adhesion gradients are not necessary, since the main parasite properties, such as stochastic motion and realistic alignment times, can be reproduced well by the bond-based model without adhesion gradients.

In conclusion, our model suggests that the parasite alignment can be explained by the passive compliance hypothesis [17, 20], such that no additional active mechanisms or processes are necessary. Of course, this does not eliminate the possible existence of some active mechanisms, which may participate in the alignment process. Another limitation of many studies is that the parasite alignment is investigated under static (no flow) conditions, whereas *in vivo*, parasite alignment and invasion occur under a variety of blood flow conditions, including different flow stresses and flow-induced RBC deformations [43]. Further experiments are needed to investigate RBC-parasite interactions for realistic blood-flow scenarios. The bond-based model proposed here is expected to be useful for the quantification of such experimental studies and for a better understanding of RBC-parasite adhesion under blood flow conditions.

## IV. METHODS & MODELS

### A. Red blood cell model

The total potential energy of the RBC model is given by [23, 24]

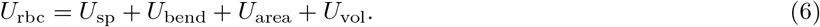

Here, the term *U*_sp_ represents the elasticity of spectrin network, which is attached to the back side of the lipid membrane. *U*_bend_ models the resistance of the lipid bilayer to bending. *U*_area_ and *U*_vol_ constrain the area and volume of RBC membrane, mimicking incompressibility of the lipid bilayer and the cytosol, respectively.

The elastic energy term *U*_sp_ is given by

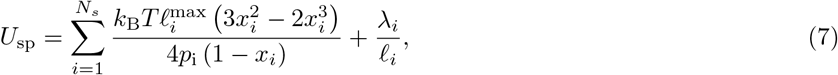

where the first term is the attractive worm-like chain potential, while the second term corresponds to a repulsive potential with a strength λ_*i*_. Furthermore, *ℓ_i_* is the length of the i-th spring, *p_i_* is the persistence length, 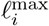 is the maximum extension, and 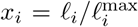. The stress-free state of the elastic network is considered to be a biconcave RBC shape, such that initial lengths in the triangulation of this shape define equilibrium spring lengths 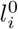. For a regular hexagonal network, its shear modulus *μ* can be derived in terms of model parameters as [23, 24]

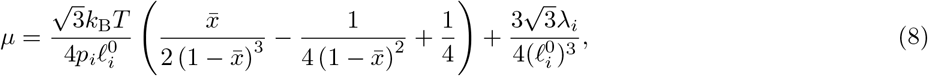

where 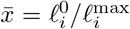 is a constant for all i. Thus, for given values of μ, 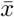, and 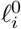, individual spring parameters *p_i_* and λ_*i*_ are calculated by using Eq. (8) and the force balance 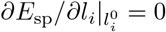 for each spring.

The bending energy of the membrane is expressed as [22, 44]

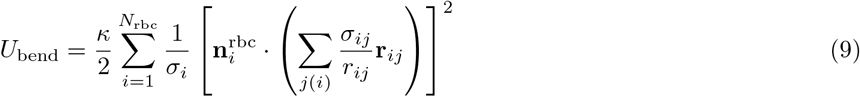

where *κ* is the bending modulus, 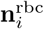 is a unit normal of the membrane at vertex *i, σ_i_* = (∑_*j*(*i*)_ *σ_ij_r_ij_*)/4 is the area of dual cell of vertex *i*, and *σ_ij_* = *r_ij_*[cot(*θ*_1_) + cot(*θ*_2_)]/2 is the length of the bond in dual lattice, with the two angles *θ*_1_ and *θ*_2_ opposite to the shared bond **r**_*ij*_.

The last two terms in Eq. (6),

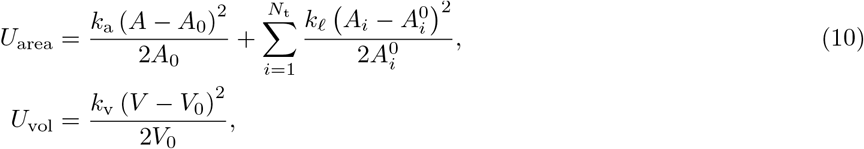

constrain surface area and volume of the RBC [23, 24], where *k*_a_and *k_ℓ_* control the total surface area *A* and local areas *A_i_* of each triangle to be close to desired total area *A*_0_ and local areas 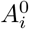, respectively. The coefficient *k*_v_ controls the total volume *V* of the cell. The values of these coefficients are chosen large enough such that the area and volume fluctuate within 1% of the desired values.

The elasticity of a healthy RBC is characterized by the shear modulus *μ* ≈ 4.8 μN m^−1^, which corresponds to the Young’s modulus *Y* ≈ 18.9 μN m^−1^for a nearly incompressible membrane. These values are employed in all simulations unless stated otherwise. The described membrane model has been shown to accurately capture RBC mechanics [23, 24] and membrane fluctuations [45].

### B. RBC-parasite adhesion interaction

Interaction between parasite and RBC membrane has two components. The first part imposes excluded-volume interactions between the RBC and merozoite (i.e. no overlap between them), using the purely repulsive part of the Lennard-Jones (LJ) potential

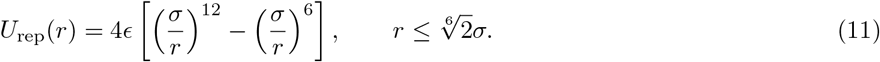

This potential acts between every pair of RBC and parasite vertices separated by a distance *r* = |**r**_rbc_ − **r**_para_| that is smaller than 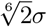. Here, *ϵ* represents the strength of the interaction and *σ* is the characteristic length scale of repulsion.

The attractive part of RBC-parasite interaction is modeled by a reversible two-state bond model. Bonds can form between the vertices at RBC membrane and merozoite surface, while existing bonds can also dissociate. These bonds represent RBC-parasite adhesion through existing agonists at the surface of these cells and can be of two different types:

i. long bonds with a maximum extension 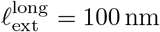,
ii. short bonds with a maximum extension 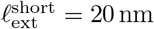,

which is motivated by electron microscopy observations of RBC-merozoite adhesion [5]. Both bond types are modeled by harmonic springs with the potential energy given by

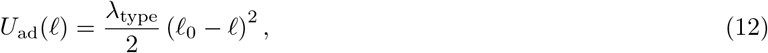

where λ_type_ is the spring constant of either long or short bond type and *ℓ*_0_ is the equilibrium length. To model the dynamic two-state interaction, <u>constant</u> (i.e. length independent) on- and off-rates (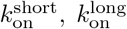, and *k*_off_) are chosen, in order to simplify the model and reduce the number of parameters. Furthermore, the off-rate for both bond types is selected to be same. Note that this model can easily be extended to length-dependent rates. To implement the different bond types, each vertex at the parasite surface can form either a long or a short bond. The choice of vertices that form long or short bonds is made randomly for fixed bond densities. To avoid possible artifacts of a single discrete bond distribution, each independent simulation assumes a different random choice of bonds with their respective densities kept constant. Bonds between the vertices at the RBC and parasite surfaces can form if the distance between two vertices is smaller than the corresponding cut-off distances 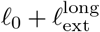 and 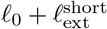, which remain the same in all simulations. Here, 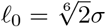 corresponds to the length of the excluded-volume LJ interactions between the vertices of RBC and parasite, whose choice is defined by a characteristic discretization length of the RBC membrane. Note that only a single bond is allowed at each vertex for the both bond types.

### C. Hydrodynamic interactions

Hydrodynamic interactions are modeled using the dissipative particle dynamics (DPD) method [27, 28], where fluid is represented by a collection of particles interacting through three types of pairwise forces: conservative 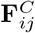, dissipative 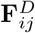, and random 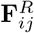 forces. The total force between particles *i* and *j* is given by

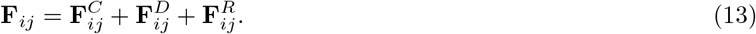

The conservative force models fluid compressibility, whereas the dissipative and random forces maintain a desired temperature of the system. The dissipative force also gives rise to fluid viscosity, which is generally measured in DPD by simulating a reversible-Poiseuille flow [46, 47]. The DPD interactions are implemented only between the pairs of fluid-fluid, fluid-RBC, and fluid-parasite particles. DPD interaction parameters are selected such that they impose no-slip boundary condition at RBC and parasite surfaces [17, 23].

### D. Simulation setup

Simulation domain with a size of 7.7*D*_0_ × 3.1*D*_0_ × 3.1*D*_0_ contains both RBC and parasite suspended in a DPD fluid, where 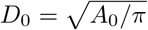 is the effective RBC diameter. Periodic boundary conditions are imposed in all directions. Initially, the parasite is placed close enough to the RBC membrane, so that the interaction between them is immediately possible. The initial parasite orientation is with its apex directed away from the membrane to mimic least favorable attachment configuration.

The main simulation parameters are shown in Table I, both in simulation and physical units. To compare simulation units to physical units, a basic length scale is defined as the effective RBC diameter *D*_0_, an energy scale as *k*_B_*T*, and a time scale as RBC membrane relaxation time 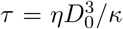. For average properties of a healthy RBC, the effective diameter is *D*_0_ ≃ 6.5 μm with *A*_0_ = 133.5 μm^2^ and the relaxation time becomes *τ* ≈ 0.92 s for the bending modulus *κ* = 3 × 10^−19^ J and plasma viscosity *η* = 1 mPa s. All simulations are performed on the supercomputer JURECA [32] at the Jülich Supercomputing Centre, Forschungszentrum Jülich.

### E. Monte-Carlo sampling of alignment times

One of the main foci of our study is to obtain distributions of parasite alignment times for various conditions, which requires a large number of simulations of merozoite alignment. In order to significantly reduce the computational effort, Monte-Carlo (MC) sampling of alignment times, which is guided by direct DPD simulations of RBC-parasite adhesion, is employed. The MC sampling is based on a two-dimensional probability map (see e.g. Fig. 4(a)), which characterizes parasite orientation at the membrane surface through the distance *d*_apex_between the parasite apex and membrane and merozoite alignment angle *θ* (see Fig. 3(a) for definitions of *d*_apex_ and *θ*). To construct such a probability map, possible *d*_apex_ and *θ* values are binned into a number of orientation states 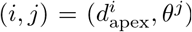, and the probability *P*(*i, j*) of each state is computed from at least 10 long DPD simulations of RBC-parasite adhesion. We have verified that 10 independent DPD simulations are enough to reliably compute a probability map through its convergence with the number of DPD simulations. In the MC algorithm, changes in parasite orientation are modeled by transitions between different states, using the Metropolis algorithm. Thus, the transition from a state (*i, j*) to one of the neighboring states (*i* +1, *j*), (*i* − 1, *j*), (*i, j* + 1) or (*i, j* − 1) is selected randomly with a probability of 1/4, and this move is accepted if *ξ < P*(new state)/*P*(*i, j*), where *ξ* is a random number drawn from a uniform distribution in the interval [0, 1]. In summary, the MC sampling algorithm is performed as follows

1. Initial parasite orientation is selected randomly by choosing a state 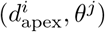, which has a non-zero probability.
2. Transitions between the neighboring states are modeled according to the Metropolis algorithm described above.
3. MC procedure is stopped whenever pre-defined alignment criteria are reached, and the number of MC steps is interpreted as alignment time.

Note that the MC sampling algorithm fulfills detailed balance, but does not account for hydrodynamic interactions. However, it is a fast and efficient way to sample the distribution of parasite alignment times.

## AUTHOR CONTRIBUTIONS

S.H. and A.K.D performed all the simulations and analyzed the computational results; G.G. and D.A.F. designed the research project; all authors interpreted the results and wrote the manuscript.

## ACKNOWLEDGMENTS

We would like to express our gratitude to Virgilio L. Lew and Pietro Cicuta from the University of Cambridge for insightful discussions. Sebastian Hillringhaus acknowledges support by the International Helmholtz Research School of Biophysics and Soft Matter (IHRS BioSoft). We gratefully acknowledge the computing time granted through JARA-HPC on the supercomputer JURECA at Forschungszentrum Jülich.

