## Supplementary material for "Stochastic bond dynamics facilitates alignment of malaria parasite at erythrocyte membrane upon invasion": Movie captions

### Supplementary information for
### "Stochastic bond dynamics facilitates alignment of malaria parasite at erythrocyte membrane upon invasion"

Sebastian Hillringhaus, Anil K. Dasanna, Gerhard Gompper, and Dmitry A. Fedosov

*Theoretical Soft Matter and Biophysics,*

*Institute of Complex Systems and Institute for Advanced Simulation,*

*Forschungszentrum Jülich, 52425 Jülich, Germany*

#### SUPPLEMENTARY MOVIES

1) **Movie S1**: Parasite motion at the membrane of a deformable RBC for the reference RBC-parasite interactions from Table 2 of the main text. $k_{\rm off}/k_{\rm on}^{\rm long} = 2$.

2) **Movie S2**: Parasite adhesion and dynamics on a deformable RBC for a reduced off-rate $k_{\rm off}$. $k_{\rm off}/k_{\rm on}^{\rm long} = 1$.

3) **Movie S3**: Parasite dynamics at the surface of a rigid RBC for the reference RBC-parasite interactions from Table 2 of the main text. $k_{\rm off}/k_{\rm on}^{\rm long} = 2$.
